# Effects of osteogenic ambulatory mechanical stimulation on early stages of BMP-2 mediated bone repair

**DOI:** 10.1101/2020.07.28.225870

**Authors:** Brett S. Klosterhoff, Casey E. Vantucci, Jarred Kaiser, Keat Ghee Ong, Levi B. Wood, Jeffrey A. Weiss, Robert E. Guldberg, Nick J. Willett

## Abstract

Mechanical loading of bone defects through rehabilitation is a promising therapeutic approach to stimulate repair and reduce the risk of non-union; however, little is known about how therapeutic mechanical stimuli modulate early stages of repair before mineralized bone formation. In a previous study, we established an osteogenic mechanical loading protocol using early ambulatory rehabilitation and a compliant, load-sharing fixator in a rat model of BMP-2 mediated bone defect repair. The objective of this study was to investigate the early effects of osteogenic loading on cytokine expression, tissue composition, and angiogenesis during the first 3 weeks of repair in this model. Using a wireless implantable strain sensor for local measurements of mechanical boundary conditions, finite element simulations showed that osteogenic mechanical loading increased mean compressive strain in defect soft tissue during rehabilitative ambulation at 1 week (load-sharing: −1.54 ± 0.17% vs. load-shielded: −0.76 ± 0.06%), and that strain was amplified in remaining soft tissue regions at 3 weeks as mineralization progressed (load-sharing: −1.89 ± 0.35% vs. load-shielded: −1.38 ± 0.35%). Multivariate analysis of multiplex cytokine arrays revealed that loading significantly altered cytokine expression profiles in the defect tissue at 2 weeks compared to load-shielded defects. Specifically, loading reduced VEGF and increased CXCL5 (LIX) levels. Subsequently, vascular volume in loaded defects was reduced relative to load-shielded defects but similar to intact bone at 3 weeks. Endochondral bone repair was also observed histologically in loaded defects only at 3 weeks. Together, these results demonstrate that moderate ambulatory strains previously shown to stimulate functional bone regeneration significantly alter early angiogenic and cytokine signaling and may promote endochondral ossification in large segmental bone defects.

**Authors’ Contributions:** B.S.K., N.J.W., and R.E.G. designed the research and performed surgeries; B.S.K., C.E.V., and J.K. performed experiments; B.S.K., C.E.V., J.K., and L.B.W., analyzed data; B.S.K., C.E.V., N.J.W., and R.E.G. wrote the manuscript; All authors interpreted data, critically edited, and have read and approved the final manuscript.

## Introduction

Bone is a dynamic tissue whose development and maintenance are heavily influenced by biomechanical stimuli exerted by load-bearing activities. After fracture, bone possesses a significant intrinsic capacity to regenerate injured tissue and restore critical biomechanical functions. However, approximately 5% of the millions of skeletal fractures that occur each year result in non-unions, where the bone defect does not heal effectively from the initial treatment and remains mechanically unstable^1^. The associated prolonged disability and multiple surgeries endured by such patients is a substantial healthcare burden^2^. Bone healing outcomes are regulated in part by the magnitude of mechanical stimulation exerted on the healing tissue during repair, where a moderate degree of mechanical stimulation can enhance bone repair, but excessive loading can result in hypertrophic non-union^3–5^.

Given the mechanosensitive nature of bone repair, the integration of rehabilitative exercise and natural load-bearing activity into treatment regimens is an established approach to stimulate bone repair after injury^5–8^. However, the majority of prior data describing therapeutic mechanical loading is predicated on relatively simple transverse fracture healing models^4,9,10^. There is little data that describes how vital biological mechanisms of bone repair prior to mineralization are modulated by mechanical loading in large segmental bone defects commonly treated with osteogenic factors including autologous bone graft or bone morphogenetic protein 2 (BMP-2). Consequently, mechanical loading is typically only implemented at later time points in segmental bone defect repair after mineralized bridging^11,12^.

Initially after a fracture, a diverse population of myeloid and lymphoid lineage immune cells rapidly infiltrate the injury site to clear pathogens and necrotic debris, remodel the extracellular matrix (ECM), and critically, to secrete cytokines to recruit mesenchymal and endothelial progenitors^13,14^. Increasing evidence has demonstrated that immune cells functionally respond to mechanical cues in a variety of disease and repair contexts^15,16^. Monocytes respond to fluid shear and compressive stresses by increasing production of pro-inflammatory cytokines in agarose gel cultures^17,18^. As immune cells resolve acute inflammatory signals, they secrete angiogenic cues. Effective bone regeneration is reliant on sufficient revascularization of the defect region via angiogenesis initiating from nearby vasculature^19^. These nascent vascular networks are also known to be mechanosensitive. Growth and alignment of microvessels in three-dimensional cultures are modulated by the application of tensile strain or dynamic compression^20,21^. The aforementioned *in vitro* results provide preliminary evidence that cytokine signaling and angiogenesis are mechanosensitive; however, *in vitro* systems do not recapitulate the cellular and molecular complexity of the regenerative niche *in vivo*. Several animal studies have demonstrated that excessive tissue loading caused by unstable bone defect fixation leads to delayed bone repair or non-union. In these impaired healing environments, there is an inhibition of angiogenic gene expression and severe abrogation of defect revascularization^22–24^. However, the effects of osteogenic and potentially therapeutic mechanical loading regimes on the biology of early stage segmental bone repair remain poorly understood.

We have previously developed a novel implantable strain sensor platform and measured strains during rehabilitation in critically-sized rat segmental defect model treated with a minimal healing dose of BMP-2^25,26^. We observed that controlled mechanical stimulation via early ambulatory loading 1 week after injury significantly enhanced bone regeneration^26^. Furthermore, we observed a significant positive correlation between strain magnitudes at 1 week with long term healing outcomes, suggesting potent magnitude-dependent effects of mechanical stimulation on early stages of bone repair. However, the underlying early-stage biological effects preceding enhanced bone formation have not been thoroughly investigated.

In this study, our overarching objective was to investigate how key aspects of the early stage regenerative response, including cytokine expression, tissue composition, and angiogenesis are modulated by the application of osteogenic mechanical strain during BMP-2 mediated bone regeneration *in vivo*. In prior studies using this pre-clinical rat segmental defect model, we have demonstrated that vascular volume under stable fixation peaks at 2 weeks and appreciable mineralization begins to occur at 3 weeks^22,27^. Thereafter, excess vessels are progressively pruned while remaining vessels mature via dilation and arteriogenesis^27,28^. Therefore, in this study, we examined cytokine expression and tissue structure during peak vascularization (2 weeks). The cytokine expression findings at 2 weeks prompted further investigation of previously collected data in this animal model to better elucidate the tissue composition and status of vascularization at the onset of mineralization (3 weeks)^26^. Given that prior research has demonstrated that excessive loading dramatically increases cytokine signaling^23^, we hypothesized that osteogenic mechanical stimulation would moderately elevate aspects of the coordinated cytokine response to support sufficient vascularization and endochondral bone repair.

## Materials and Methods

### Surgical procedure

All animal procedures were approved by the Georgia Institute of Technology IACUC (Protocol A17034). Two animal studies were conducted using identical injury, treatment, and rehabilitation procedures described below. The first study concluded 2 weeks post-surgery with multiplex cytokine and histological analyses of the newly formed tissue in the bone defect. The findings of the 2 week analyses prompted us to conduct new analyses utilizing mechanical strain data and tissue samples collected during a previous animal study in this model which concluded at 3 weeks with terminal microcomputed tomography (microCT) angiography. Therefore, microCT-based finite element models, histological samples, and vascular volumes from this previous study were generated and analyzed to better understand the progression of the mechanical environment as well as the tissue composition and vascular volume at the onset of mineralization in this model^26^.

Similar to previously described procedures, a unilateral 6 mm segmental defect was surgically created in the left femur of 15 week old female CD Sprague-Dawley rats (n=24, Charles River Labs)^29^. Anesthesia was maintained by isoflurane inhalation and analgesia was provided by a pre-operative subcutaneous injection of sustained-release buprenorphine. Before creating the defect with an oscillating saw, femurs were stabilized by affixing a custom-machined radiolucent internal fixator plate comprised of either polysulfone (PSU, McMaster-Carr), or ultra-high molecular weight polyethylene (UHMWPE, Quadrant) using four stainless steel screws. UHMWPE fixators were 40% more compliant than PSU (flexural stiffness: PSU = 232 ± 20 N/mm; UHMWPE = 145 ± 11 N/mm)^26^. Fixators in the 3 week vascularization study contained integrated wireless strain sensors that provided real-time non-invasive measurements of mechanical strain across the bone defect during ambulatory activity, as previously described^25,26^. The sensors transmitted wireless measurements of axial strain across the bone defect fixator in real-time during ambulation via Bluetooth. Fixators in the 2 week defect tissue cytokine analysis study were stabilized by identical fixators without the integrated strain sensors. Defects in both studies were acutely treated with a minimal healing dose of recombinant human bone morphogenetic protein 2 (BMP-2) delivered via a hybrid RGD-alginate/PCL biomaterial platform described in detail previously^30,31^. Briefly, treatment consisted of installing an electrospun polycaprolactone (PCL, Sigma-Aldrich) tube across the defect gap, then injecting 120 μL alginate hydrogel laden with 2 μg BMP-2 (Pfizer) inside the tube. Fixators were allocated randomly. After either 2 or 3 weeks, animals were anesthetized then euthanized via CO_2_ asphyxiation.

### Treadmill walking

To exert a semi-controlled ambulatory mechanical load on the femur, animals were walked on a treadmill. Animals were trained to walk at a slow, consistent speed of 6.5 m/min 1 week prior to surgery. After surgery, animals were allowed to recover in cages for 1 week, then 10 minute walking periods were initiated at day 7 and continued twice weekly thereafter. The distance traversed during this period (65 m) roughly approximates the best available estimate of the total distance covered during one day of in-cage activity in healthy mice^32^.

### Finite element analyses

Sample-specific finite element (FE) models of the femur-fixator constructs were constructed at 1 and 3 weeks using similar methods described previously^26^. Briefly, microCT images were processed in MIMICS (Materialise) to create a representative model of proximal and distal femur segments with the fixator attached. At 1 week, defects were assumed to be a homogeneous 5 mm diameter cylinder comprised of soft tissue. For 3 week models, pre-decalcification microCT geometry of mineralized woven bone (threshold = 388 mg HA/ccm) were included within the soft tissue cylinder of each femur sample. Volumetric meshes of quadratic tetrahedral elements were constructed. Mesh resolution convergence was previously demonstrated at approximately 1.5×10^6^ equations^26^. All materials were constitutively modelled as neo-hookean solids and previously reported experimentally validated mechanical properties were assigned (Table S1)^26,33^.

In each model, animal-specific boundary conditions were defined based on axial strain measurements by the wireless strain sensor at the corresponding 1 or 3 week treadmill period. Similar to previously reported simulations, boundary conditions were applied to replicate *in vivo* loading^34^; the distal end of the femur was fixed in all directions and combined axial compression and bending pressures were applied to the proximal surface. All FE simulations were conducted in FEBio (version 2.8.5) and results were evaluated in PostView^35^. To capture inter-animal variability of the complete data set while maintaining computational efficiency, a sub-set of animals (n=5 from each fixator group) were evaluated by longitudinal FE analyses at both 1 and 3 weeks, creating 20 FE simulations in total. The distribution of strain magnitude and woven bone volume were identical between the sub-set of animals and the complete experimental group, confirming that the sub-set was an accurate depiction of the full data set (Fig. S1).

### Analyses of defect tissue cytokines

In the 2-week study, animals were euthanized 24 hours after the final week 2 treadmill period. Defect tissue inside and adhered to the outside of the PCL tube was immediately and carefully harvested. Upon dissection, two samples were excluded from analysis due to evidence of tissue abscess near the fixator indicative of infection. Individual tissue samples were flash frozen in liquid nitrogen and then stored at −80° C for subsequent Luminex multiplexed immunoassays. Samples were thawed and homogenized in RIPA lysis buffer (Thermo Fisher) supplemented with 1X Halt protease inhibitor cocktail (Thermo Fisher). Lysates were centrifuged at 13,000 g for 15 minutes and supernatants were transferred to new tubes and stored at −80° C. A panel of 27 cytokines were quantified using a Milliplex MAP Rat Cyotkine/Chemokine Magnetic Bead Panel according to the manufacturer’s instructions and then read using a MAGPIX instrument (Luminex). Cytokine concentration was normalized to total protein content quantified using a bicinchoninic acid assay (BCA assay, Thermo Fisher). The final sample sizes for each group were: UHMWPE = 9 and PSU = 11.

### Histology

Representative femur samples from each study end point, at weeks 2 and 3 respectively, were reserved for histological analysis based on digital radiographs acquired immediately before euthanasia. Femora were fixed in 10% neutral-buffered formalin for 48 hrs at 4° C and switched to PBS after. HistoTox Labs completed decalcification, paraffin processing, and staining. Midsagittal 5 μm thick sections were stained with Hematoxylin & Eosin (H&E), Safranin-O/Fast Green, or Picro Sirius Red.

### MicroCT angiography

Vascular volume within and around the segmental bone defect in the 3 week animal study was quantified using microcomputed tomography (microCT) angiography. Briefly, vasculature was perfused successively with 0.9% saline, 0.4% papaverine hydrochloride vasodilator, 0.9% saline, 10% neutral buffered formalin (NBF), 0.9% saline, and radiopaque contrast agent^36^ (2:1; Microfil MV-22, FlowTech Inc.), then allowed to polymerize at 4° C overnight. Dissected femora were scanned using microCT with 15 μm voxels, 55 kVp, 145 μA, and 300 ms integration time. Samples were then decalcified for 10 days under light agitation in a mixture of formic acid and citrate (Newcomers Supply) changed daily. Samples were then scanned with microCT a second time. Vascular volume was quantified within 3 different 4.14 mm long cylindrical volumes of interest (VOI’s): (1) the “Defect VOI” – a 5 mm diameter cylinder within the PCL mesh tube spanning the defect; (2) the “Total VOI” – a 7 mm diameter cylinder encompassing the Defect VOI and including surrounding vasculature adjacent to the PCL mesh; and (3) the “Surrounding VOI” - the annulus created by subtracting the Defect VOI from the Total VOI.

### Statistical analyses

Data are displayed as mean ± standard error of the mean or as box plots showing 25th and 75th percentiles, with whiskers extending to minimum and maximum values, unless otherwise noted in the figure heading. Differences were assessed using t-test. Welch’s t-test or Mann-Whitney U test and Kruskal-Wallis test was used in cases of unequal variances or non-normal distributions, respectively. Two-way ANOVA with Tukey’s or Sidak’s pairwise comparisons were used to compare groups for longitudinal FE analyses. Significance was determined using p < 0.05. Statistical tests were performed by GraphPad Prism 8.

Univariate and multivariate analysis of cytokine expression profiles within the defect were conducted to evaluate differences between fixator stiffness groups, as described previously^37^. Briefly, univariate t-tests were performed comparing the normalized median fluorescence intensity values of individual cytokines. Discriminant partial least squares regression (D-PLSR) was performed using the PLS MATLAB (Mathworks) algorithm developed by Cleiton Nunes (Mathworks File Exchange). Individual cytokine measurements were used as independent variables, and fixator stiffness was used as the discrete regression variable. Multi-dimensional latent variables (LV’s) were defined in three dimensions, and an orthogonal rotation was implemented to assess separation by fixator stiffness in the LV1-LV2 plane. Standard deviation of the contribution of each cytokine to LV1 were computed using leave-one-out cross validation without replacement.

## Results

### Local defect tissue strain was elevated by load-sharing fixation and localized to non-mineralized soft tissue regions

Specimen-specific longitudinal FE analyses revealed elevated 3^rd^ principal strain magnitude (the maximum local compressive strain) during gait within defects stabilized by compliant, load-sharing UHMWPE fixation at 1 week (Fig. 1 & Fig. 2A & B; UHMWPE: −1.54 ± 0.17%; PSU: −0.76 ± 0.06%, p = 0.077). Between 1 and 3 weeks, woven bone extending from the intact bone ends was formed within the defects. The deposition of relatively stiff woven bone reduced the proportion of soft tissue within the defect. Consequently, the magnitude of 3^rd^ principal strain within soft tissue increased between weeks 1 and 3 (Fig. 2A & C Wk 3: UHMWPE: −1.89 ± 0.35%; PSU: −1.38 ± 0.35%, *p = 0.04 Wk 1 vs. 3). As indicated by representative FE model cross-sections, the magnitude of strain was relatively heterogeneous throughout the defect, reaching maxima of 3-6% in some regions under load-sharing fixation at 1 week and increasing to 5-10% in both fixation groups at 3 weeks (Fig. 1 & Fig. 2B & C). In newly formed woven bone, mean 3^rd^ principal strain magnitude was similar regardless of fixator stiffness (Fig. 2D & E; UHMWPE: −0.47 ± 0.08%; PSU: −0.55 ± 0.12%, n/s). The resultant axial load at the proximal femur during gait averaged 3.78 ± 0.35 bodyweight (BW) overall (Fig. 2F). Axial load did not change between 1 and 3 weeks and was not affected by fixation stiffness. Axial loads had high variance from animal to animal (Min: 0.87 BW; Max: 7.2 BW), and there was a significant effect of animal over time (p < 0.05), suggesting heterogeneity in hindlimb utilization on a per animal basis as each animal recovered from surgery.

**Fig. 1.**
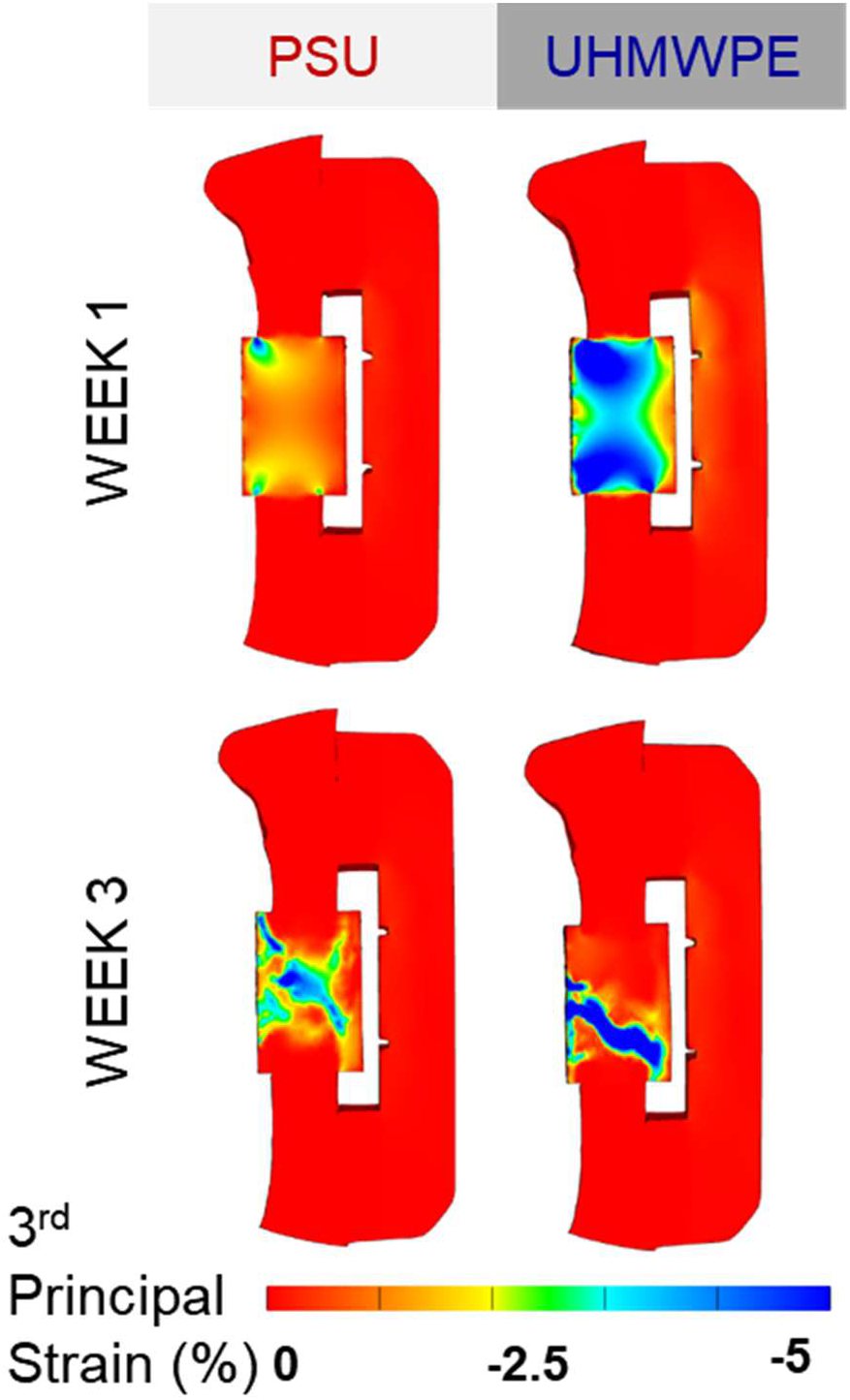
Representative image-based finite element model cross-sections demonstrate elevated strain magnitudes within UHMWPE-stabilized defects at 1 week. Regions of elevated strain localized to non-mineralized soft tissue regions at 3 weeks.

**Fig. 2.**
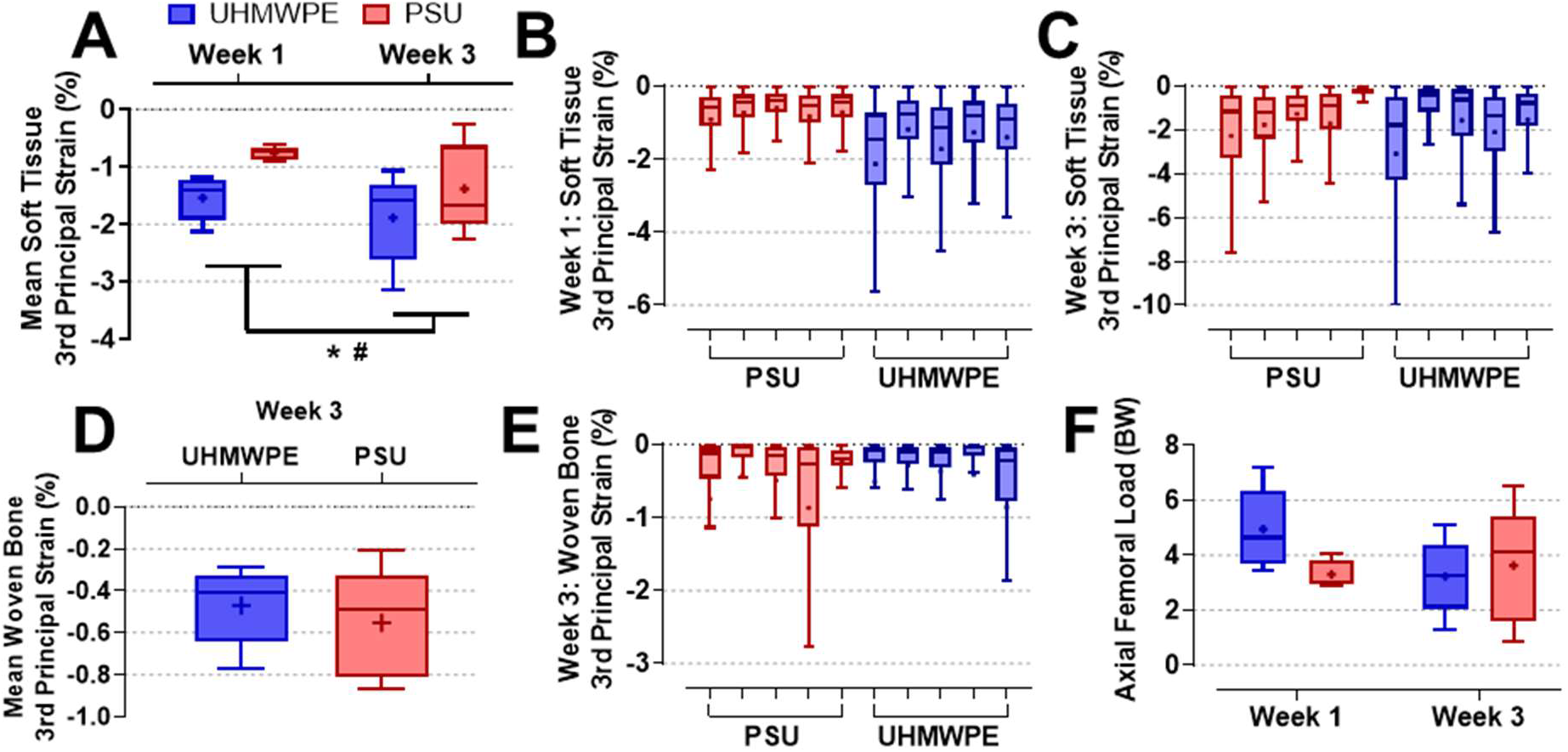
(A) Mean 3rd principal strain of non-mineralized soft tissue within the defect increased between 1 and 3 weeks due to the increased proportion of much stiffer woven bone as healing progressed. n=5. *p<0.05 Wk 1 vs. 3, #p=0.077 UHMWPE vs. PSU via Two-way RM ANOVA. Soft tissue strain distribution throughout the defect within each of 5 experimental samples was spatially heterogeneous at (B) 1 and (C) 3 weeks. In all box plots, the mean is denoted by a dot and whiskers are defined using the Tukey method. (D) Mean 3rd principal strain magnitude within immature woven bone at 3 weeks averaged approximately 0.5% regardless of fixator stiffness. n=5. n/s via t-test. (E) Woven bone tissue strain was similarly heterogeneous throughout the defect in each experimental sample. (F) Peak femoral loads were variable between animals, averaging approximately four-fold body weight (BW) regardless of fixator stiffness or time point. n=5. n/s via Two-way RM ANOVA.

### Increased strain magnitude altered cytokine expression profile within bone defects at 2 weeks

Defect tissue lysate was analyzed for an array of cytokines. To distinguish multivariate cytokine expression profiles differentially regulated by defect tissue strain, discriminant partial least squares regression (D-PLSR) was performed^38^. D-PLSR revealed that fixation plate stiffness groups primarily separated along the LV1 axis (Fig. 3A). Defect samples stabilized by stiffer, load-shielding PSU fixators scored significantly higher along LV1 relative to samples stabilized by more compliant, load-sharing UHMWPE fixators (Fig. 3B). The LV1 axis represents a combination of cytokines that have been reduced in dimensionality in order to maximally separate the data based on fixator stiffness. Based on the LV1 loading plot, we can determine which cytokines most contributed to positive and negative LV1 scores, which were associated with load-shielding and load-sharing fixation, respectively (Fig. 3C). VEGF and IL-4 were associated with load-shielding along LV1, whereas expression of LIX (CXCL5), IL-1β, and RANTES (CCL5) was associated with load-sharing fixation along LV1 (Fig. 4). Univariate pairwise comparisons showed that LIX (CXCL5) levels were significantly elevated by load-sharing fixation. Conversely, VEGF expression was elevated by load-shielding fixation (p=0.052).

**Fig. 3.**
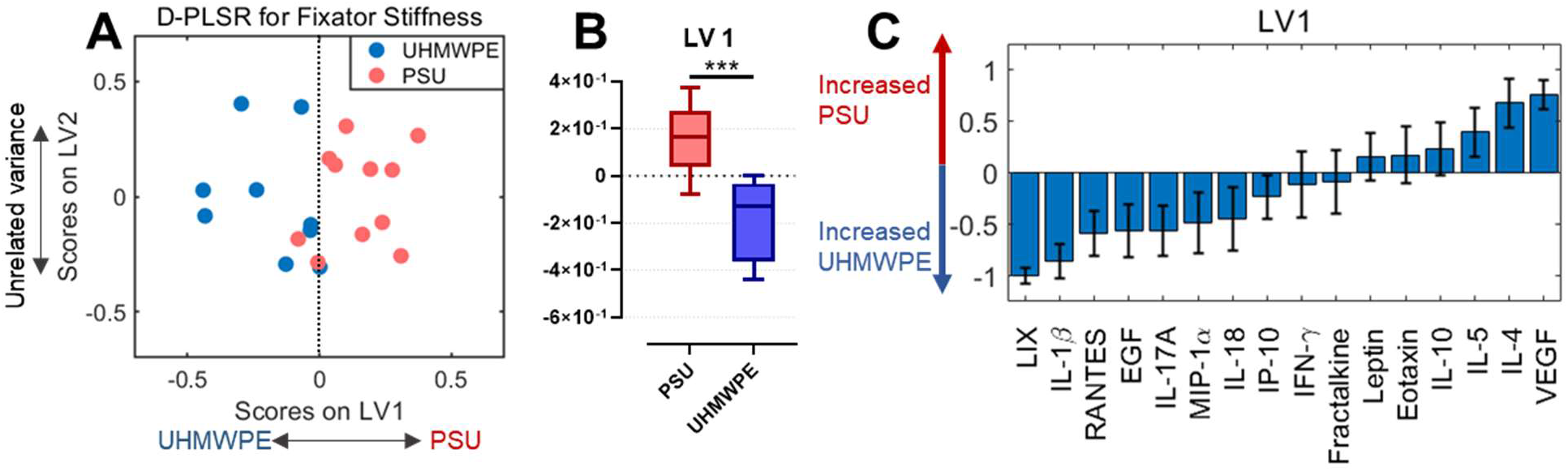
Discriminant partial least squares regression (D-PLSR) analysis of the expression of 16 cytokines in the defect tissue at 2 weeks. (A) Latent variable 1 (LV1) defines a multivariate cytokine expression profile depicted along the x-axis that separates defects stabilized by load-shielding PSU fixators to the right and load-sharing UHMWPE fixators to the left. (B) Mean LV1 score was significantly upregulated by load-shielding PSU fixation. n=9-11. ***p<0.001 via t-test. (C) Values along LV1 describe individual cytokines with elevated expression for UHMWPE fixation (negative values) or PSU (positive values). Error bars were computed using leave-one-out cross validation (mean ± SD).

**Fig. 4.**
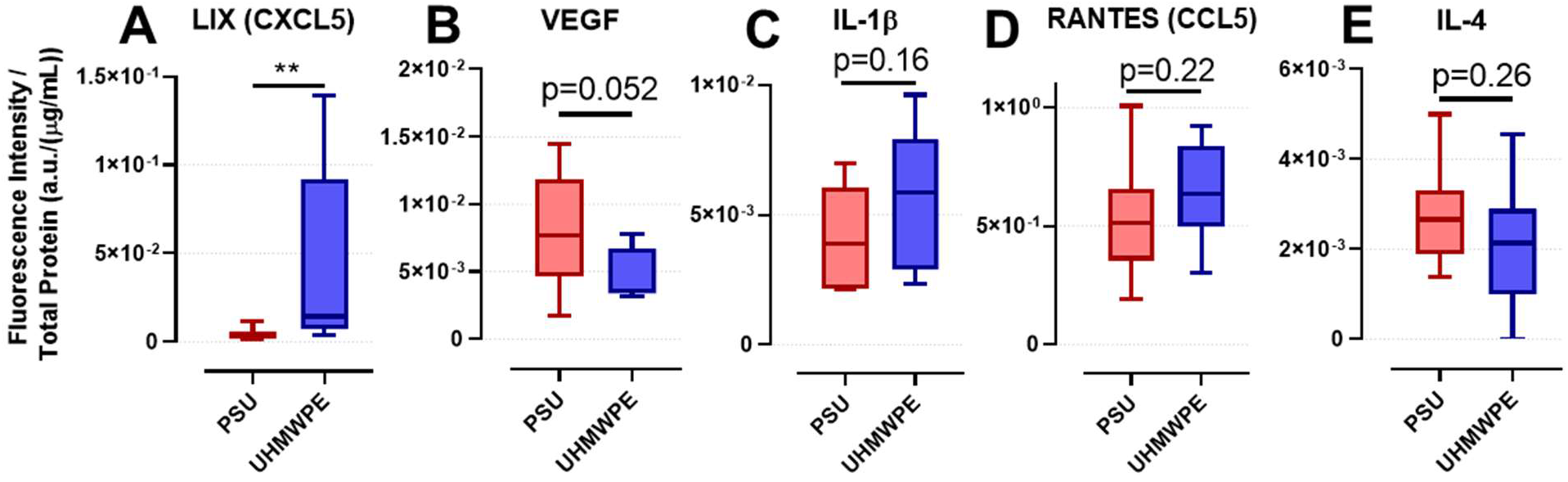
Univariate comparisons of individual cytokine expression in defect tissue at 2 weeks. LIX (CXCL5) expression was significantly increased by increased load sharing with load-sharing UHMWPE fixation. Conversely, VEGF expression was elevated with by load shielding with load-shielding PSU fixation. Significant pairwise differences were not observed in the remaining cytokines. n=9-11. **p<0.01 via t-test.

### Vascular volume within the defect was increased at 3 weeks by load-shielding PSU fixation

MicroCT angiography revealed that vascular volume within the defect was significantly increased under stiffer, load-shielding PSU fixation at 3 weeks relative to compliant, load-sharing UHMWPE fixation or unoperated contralateral femora (Fig. 5A). The vascular volume in defects stabilized by UHMWPE was similar to the contralateral. Changes in vascular volume mediated by fixation plate stiffness were only present within the central defect; there were no differences between PSU and UHMWPE fixators in the total analyzed volume or in the surrounding tissue alone (Fig. 5B & C). Regardless of fixator stiffness, vascular volume surrounding the bone defects was significantly increased relative to unoperated femora (Fig. 5C).

**Fig. 5.**
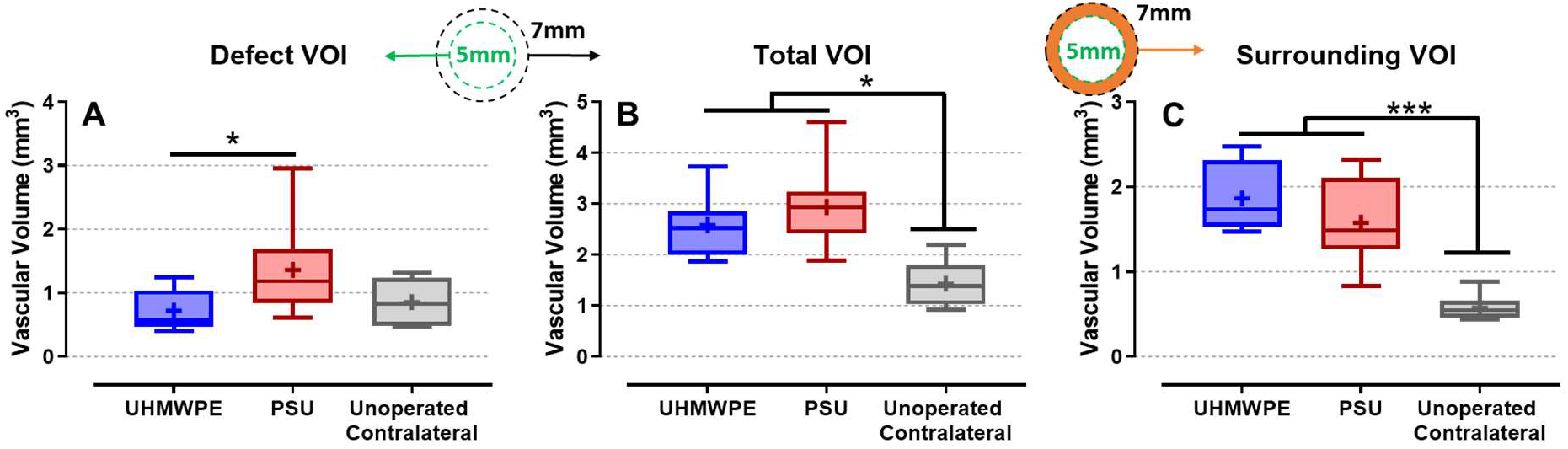
MicroCT angiography quantification of vascular volume at 3 weeks in and surrounding the bone defects. (A) Vascular volume within the central 5 mm of the defect is increased under load-shielding fixation, though load-sharing fixation remained similar to the intact femora. n=6-10. *p<0.05 via Kruskal-Wallis test with Dunn’s pairwise comparisons (B) Vascular volume throughout the defect and surrounding tissue was elevated in injured femora relative to naïve, irrespective of fixation stiffness. n=6-10. *p<0.05 via ANOVA with Tukey’s test. (C) In the surrounding tissue alone, vascular volume was similarly elevated in injured hindlimbs regardless of fixator. n=6-10. *p<0.05 via ANOVA with Tukey’s test.

### Mechanical loading promoted endochondral woven bone formation at 3 weeks

Qualitative histological analysis of defect tissue samples at 2 and 3 weeks revealed that marked changes in tissue composition occur during this relatively brief interval (Fig. 6). At 2 weeks, the defect was predominantly comprised of alginate hydrogel from the BMP-2 delivery vehicle. Irrespective of fixation stiffness, the presence of alginate declined between weeks 2 and 3, while qualitative increases in mineralized tissue were simultaneously apparent. Hypertrophic chondrocytes were evident at 3 weeks throughout defects stabilized by load-sharing UHMWPE but not in stiffer PSU, suggesting the initiation of endochondral ossification due to mechanical load sharing. Picro Sirius red staining viewed under polarized light revealed large red collagen fibers adjacent to regions of hypertrophic chondrocytes with UHMWPE, indicative of woven bone formation. Collagen in PSU-stabilized samples appeared primarily orange-yellow in color and qualitatively smaller in size. Lamellar collagen architecture was not observed in either group at these relatively early time points.

**Fig. 6.**
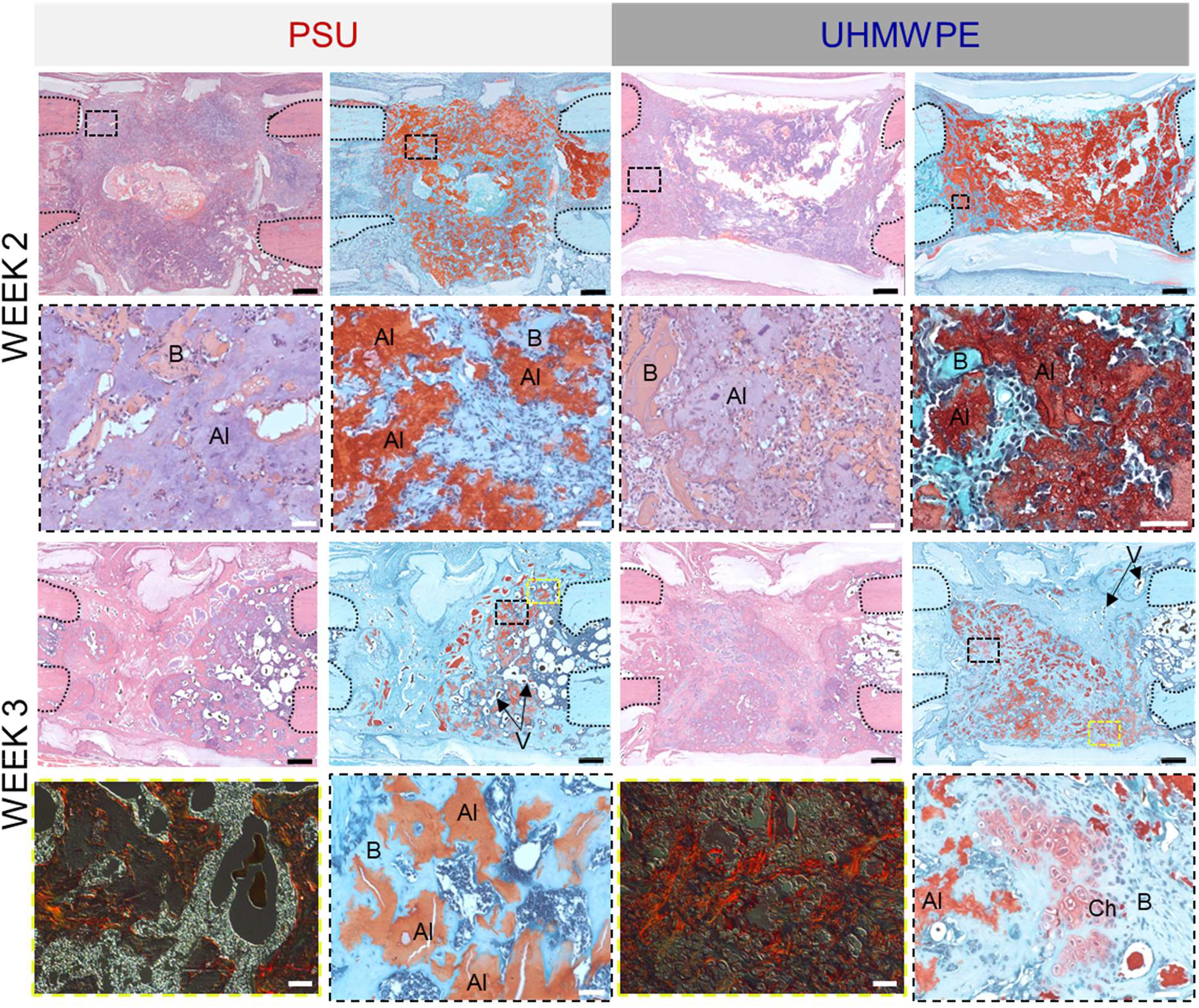
Bone defect histology at 2 and 3 weeks, including Hematoxylin & Eosin (left column for PSU and UHMWPE, except for bottom image), Safranin0O/Fast Green (right columns), and Picro Sirius red (bottom image in left columns). Alginate hydrogel remnants are denoted as “Al”, bone is denoted as “B”, chondrocytes are denoted as “Ch”, and blood vessels are indicated with arrows extending from “V”. Dotted lines at sides of images denote intact diaphyseal bone stumps. Insets are registered to full defects by color-coded dotted rectangles. Black scale bar = 500 μm. White scale bar = 50 μm.

## Discussion

Controlled mechanical loading of bone defects is a potentially potent therapeutic approach to stimulate bone repair and reduce non-union risk, but the effects of stimulatory mechanical loading on the early stages of bone repair not well understood. In particular, mechanical loading may enhance bone repair in clinically challenging defects including segmental defects created surgically after the resection of tumors, debrided comminuted traumatic fractures, or spinal fusions and corpectomies; all of which are generally treated with bone grafting or BMP-2 to supplement osteogenic capacity. The osteogenic effects of mechanical loading on bone repair after defects have been bridged with immature mineralized tissue are well described^28,39^. However, there is little data characterizing how potentially beneficial biophysical stimuli modulate the early stages of tissue repair after skeletal injury and before mineralized tissue is formed. Increasing evidence has demonstrated the fundamental role of the cytokine response in coordinating cell recruitment and angiogenesis preceding mineralization, and *in vitro* studies support that immune mediators alter cytokine secretion in response to mechanical stimuli^15–17,40–42^. Elucidating the interplay between biophysical stimulation and early stage bone repair is critical to gain a mechanistic understanding of how best to employ mechanical therapies to benefit patients at elevated risk of non-union.

In a previous study we developed a bone defect mechanical loading regimen consisting of biweekly ambulation and a load-sharing internal fixator instrumented with a wireless strain sensor; the increased mechanical stimulation permitted by the load-sharing fixator enhanced bone regeneration by 60% and nearly tripled the percent of bridged defects relative to a stiffer, load-shielding fixator. In this study, we probed how this osteogenic mechanical loading regimen altered the early stage molecular and tissueprogression of bone defects. To characterize the spatiotemporal progression of this osteogenic mechanical environment *in vivo*, FE simulations of bone defects at 1 and 3 weeks were constructed using experimental *in vivo* strain measurements as boundary conditions. Compressive strain magnitudes in the non-mineralized defect soft tissue under more compliant, load-sharing fixation averaged 1.5% and 1.9% at 1 and 3 weeks respectively. As deposition of comparatively stiff woven bone was initiated, the strain magnitude within non-mineralized tissue in the defect increased between 1 and 3 weeks as deformation localized to remaining soft tissue regions. Spatial heterogeneity in deformation was apparent at both 1 and 3 weeks, though soft tissue regions with locally elevated strain magnitudes remained below 10-15% strain—a biomechanical threshold previously implicated to cause fibrosis and hypertrophic non-union^4,43,44^. A limitation of the 3-week finite element models was that blood vessels perfused with radiopaque contrast agent could not be distinguished from woven bone on microCT, though vascular volume represented a small proportion of the thresholded tissue volume, less than 7%, compared to bone. FE simulations also revealed that animals exerted an axial load at the proximal femur of approximately 4-fold bodyweight during rehabilitative treadmill walking sessions which was unaffected by fixation stiffness or time. Considering the bone injury and iatrogenic effects of the surgery, these results are comparable to an inverse dynamics analysis by Wehner and colleagues that estimated that the peak axial load through the proximal femur in an uninjured rat was 6-fold bodyweight^34^

Changes in the tissue-level mechanical environment imparted by compliant, load-sharing fixation altered early stage cytokine expression. Multiplex cytokine analysis revealed that VEGF levels in the defect tissue lysate at 2 weeks were downregulated by compliant fixation. Previous studies have shown reduced VEGF mRNA expression observed at 4, 7, and 21 days in a highly unstable sheep osteotomy model that delayed bone repair^24^—though a clear difference being that VEGF levels still reached a sufficient level for bone repair in the study we present here. These data suggest that exogenously delivered BMP in a load shielded environment may induce VEGF exceeding the necessary levels, while unstable fixation may attenuate VEGF levels below the necessary threshold for bone healing. In addition to VEGF, expression profiles of immune cytokines and chemokines were also differentially regulated by strain magnitude. Multivariate expression profile analysis indicated LIX (CXCL5), IL-1β, and RANTES (CCL5), cytokines that exert diverse effects on angiogenesis, inflammation, and cell differentiation, and tissue repair, were positively associated with mechanical loading. LIX (CXCL5) expression has been shown to enhance angiogenesis, inhibit osteoclastogenesis, and can be secreted by mesenchymal progenitors via non-canonical WNT signaling as well as by macrophages^45–48^. IL-1β is classically defined as a pro-inflammatory mediator after tissue injury^49^. However, even after the resolution of acute inflammation, IL-1β signaling has also been shown to play subtle roles stimulating various phases of tissue regeneration, including VEGF production, endochondral ossification, and bone remodeling^50–54^. Similarly, RANTES (CCL5) can promote VEGF production and angiogenesis and osteogenic differentiation of mesenchymal stem cells^48,55–57^. There is little comparative data describing the acute cytokine response to osteogenic mechanical loading *in vivo*. Excessive mechanical loading designed to impair healing in a mechanically unstable sheep osteotomy model resulted in elevated cytotoxic T cells and reduced expression of the endothelial cell marker von Willebrand factor (vWF) 60 hours after surgery compared to stable fixation^23^. Pro-inflammatory cytokine production has also been reported in monocytes subjected to shear and compression in three-dimensional cultures. Taken together, these data suggest that mechanical loading may exert magnitude dependent effects on angiogenic and inflammatory signaling, which at low compressive magnitudes (<7%) prior to mineralization may bolster tissue repair, but at excessive magnitudes (>10%) may severely inhibit angiogenesis and impair healing^26^. However, the single time point of analysis and dynamic nature of cytokine production limit definitive conclusions on the overall time course of angiogenic and inflammatory signal production.

Vascular infiltration into the defect at 3 weeks was also significantly altered by dynamic strain magnitude. Supporting increased levels of VEGF in the defect tissue lysate at 2 weeks under stiffer, load-shielding fixation, reduced strain led to elevated vascular volume in the defect at 3 weeks. However, the relatively larger magnitude deformation with UHMWPE fixation still supported similar vascular volume to intact femora. In response to injury, angiogenic sprouting typically produces excess nascent vessels that exceed physiological demands initially, and insufficiently matured or surplus vessels are subsequently trimmed^58,59^. Prior angiography studies in the rat femoral segmental defect model demonstrated that vascular volume with load-shielded PSU fixation peaks at 2 weeks; thereafter, excess vessels are progressively pruned while remaining vessels mature via dilation and arteriogenesis^27,28^. Therefore, the 3-week time point was selected to capture the status of vascularization after the pruning and maturation process begins and the onset of mineralization in the defect. Given the single 3-week measurement of vascularization, a limitation of this study is that it cannot be explicitly determined if increased strain magnitude permitted by load-sharing UHMWPE fixation reduced or delayed angiogenic sprouting, or accelerated pruning. Regardless, vascular volume with load-sharing fixation was similar to intact bone, suggesting that the threshold of sufficient vascularization had been achieved.

Consistent with prior reports, larger magnitude strain in the range of 1-7% permitted by compliant, load-sharing fixation in the presence of BMP-2 supported endochondral ossification at 3 weeks, whereas intramembranous ossification was predominant with stiffer, load-shielding fixation^12,28^. We previously observed the presence of hypertrophic chondrocytes in this mechanical loading model at 8 weeks, at which point there was a significant 60% increase in bone formation over load-shielding fixation^26^. Together, these findings support that BMP-2 mediated skeletal repair can proceed via either intramembranous or endochondral ossification, and that the biomechanical conditions can dictate this switch. Overall, the data indicate there is a strain magnitude threshold where vascularization begins to be reduced or delayed slightly while bone repair is simultaneously improved, primarily via endochondral repair. Prior work using significantly more compliant fixation systems have demonstrated that further increases in local tissue strain exceeding 10-15% dramatically inhibits both angiogenesis and osteogenesis^4,28^. Therefore, the biomechanical environment produced by rehabilitative or mechanical therapies must be carefully monitored to be safe and efficacious.

The cytokine response is critical to coordinate angiogenesis and skeletal repair after injury. The data presented here suggest that cytokine signaling processes coordinating bone repair are mechanosensitive and that key cytokines are differentially expressed when subjected to osteogenic mechanical loading produced by rehabilitative activity. The findings of this study motivate further research to determine the downstream effects of mechanosensitive perturbations in cytokine signaling on specific progenitor cell sub-populations that mediate osteogenesis.

## Acknowledgements

Funding for this work was provided by grants from the National Institutes of Health (NIH R21 AR066322; NIH R01 AR069297) and the National Science Foundation (NSF CMMI-1400065). The content is solely the responsibility of the authors and does not necessarily represent the official views of the National Institutes of Health or National Science Foundation. This work also received support from VA (Merit) Grant RX001985 from the United States (U.S.) Department of Veterans Affairs Rehabilitation Research and Development Service. The contents do not represent the views of the U.S. Department of Veterans Affairs or the United States Government. B.S.K. received support from the Cell and Tissue Engineering NIH Biotechnology Training Grant (T32-GM008433) and the National Science Foundation Graduate Research Fellowship Program (DGE-1650044). C.E.V. received support from the National Science Foundation Graduate Research Fellowship Program (DGE-1650044). The authors gratefully acknowledge assistance and expertise provided by members of the Guldberg and Willett labs during surgeries, the core facilities at the Parker H. Petit Institute for Bioengineering and Bioscience, and the Georgia Tech Physiological Research Lab. We also gratefully acknowledge Dr. Johnna Temenoff for permitting access to the rodent treadmill, and Dr. Scott Hollister for access to image processing software.

## Supplementary Figures & Tables

**Table S1:** Finite element model mechanical properties.

| Material | E (MPa) | $\nu$ |
| --- | --- | --- |
| Intact cortical bone | $18 \times 10^3$ | 0.33 |
| Intact trabecular bone | 500 | 0.33 |
| Soft defect tissue<sup>1</sup> | 0.022 | 0.4 |
| Mineralized defect tissue<sup>2</sup> | 36.2 | 0.33 |
| PCL tube<sup>1</sup> | 1.44 | 0.37 |
| PSU fixator<sup>1</sup> | 844 | 0.37 |
| UHMWPE fixator<sup>1</sup> | 525 | 0.4 |
| Steel anchor plate | $200 \times 10^3$ | 0.3 |

<sup>1</sup>Klosterhoff, et al., Bone, 2020.
<sup>2</sup>Leong & Morgan, Integrative and comparative biology, 2009.

**Fig. S1.**
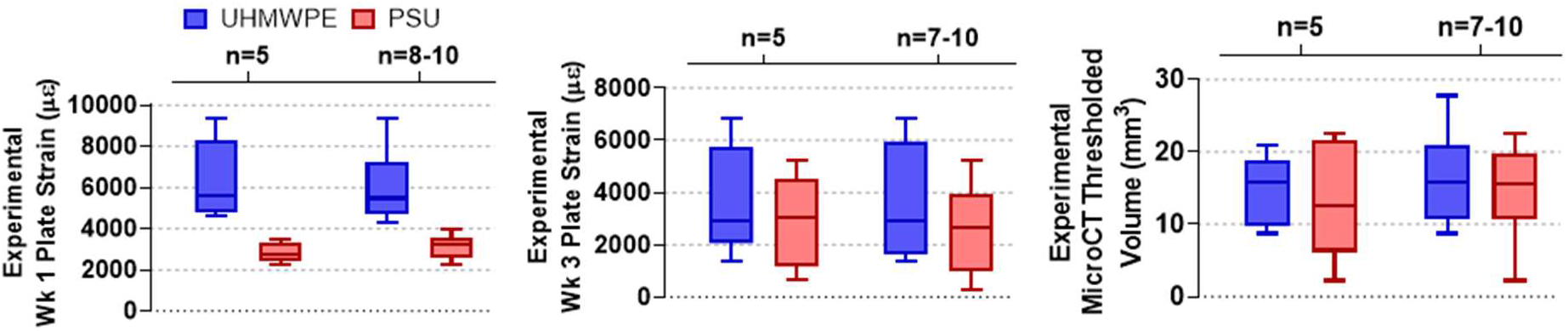
Finite element (FE) analyses were conducted on a sub-set of experimental samples representative of the complete data set. Longitudinal FE simulations were conducted on 5 experimental samples from each experimental group. Key experimental inputs for the FE simulations including magnitudes of fixation plate strain at 1 and 3 weeks (the boundary conditions), as well as the thresholded tissue volume at 3 weeks (the volume of woven bone) of the sub-sets were identical to the complete data set for each group.

